# Krüppel-like Factor (KLF) family members control expression of genes required for serous cavity and alveolar macrophage identities

**DOI:** 10.1101/2024.02.28.582578

**Authors:** Kathleen Pestal, Leianna C Slayden, Gregory M Barton

## Abstract

Tissue-resident macrophages adopt distinct gene expression profiles and exhibit functional specialization based on their tissue of residence. Recent studies have begun to define the signals and transcription factors that induce these identities. Here we describe an unexpected and specific role for the broadly expressed transcription factor Kruppel-like Factor 2 (KLF2) in the development of embryonically derived Large Cavity Macrophages (LCM) in the serous cavities. KLF2 not only directly regulates the transcription of genes previously shown to specify LCM identity, such as retinoic acid receptors and GATA6, but also is required for induction of many other transcripts that define the identity of these cells. We identify a similar role for KLF4 in regulating the identity of alveolar macrophages in the lung. These data demonstrate that broadly expressed transcription factors, such as Group 2 KLFs, can play important roles in the specification of distinct identities of tissue-resident macrophages.

**SUMMARY:** Previous studies have identified many specific regulators of macrophage development. This work reveals the requirement of members of the Group 2 KLF transcription factor family in the determination of the identity of distinct tissue-resident macrophages.

## INTRODUCTION

Tissue-resident macrophages develop from yolk sac and fetal liver progenitors during embryogenesis (Mass *et al*., 2016; Stremmel *et al*., 2018; Gomez Perdiguero *et al*., 2015). The essential macrophage growth factor M-CSF (CSF-1) induces these progenitors to differentiate into early macrophages that seed the developing tissues where they encounter additional, tissue-specific signals that drive the expression of distinct tissue phenotypes and functions (Gautier *et al*., 2012; Gosselin *et al*., 2014; Lavin *et al*., 2014). Previous studies have demonstrated the importance of such tissue-specific signals by performing macrophage transfers between tissues or tracking differentiation of monocytes as they repopulate a tissue after perturbation (Lavin *et al*., 2014; Sakai *et al*., 2019; Roberts *et al*., 2017). For some tissue-resident macrophage populations, key transcription factors have been identified that respond to tissue-specific signals and control enhancer activity and gene expression, ultimately leading to the discrete functional characteristics of each population(Sakai *et al*., 2019; Schneider *et al*., 2014; Okabe and Medzhitov, 2014; Guilliams *et al*., 2013; Buttgereit *et al*., 2016). However, for most tissue-resident macrophage populations, our understanding of the transcriptional networks required for this specification remains incomplete.

The Krüppel-like Factors (KLFs) are a family of 17 broadly expressed eukaryotic zinc finger transcription factors consisting of three subgroups (Yuce and Ozkan, 2024; Pollak *et al*., 2018). The KLFs are involved in many different aspects of biology, functioning as both repressors and activators of transcription. In several studies cataloging the different molecular identities of tissue-resident macrophages, an enrichment in Krüppel-like Factor (KLF) transcription factor binding sites was noted in regions of active and open chromatin of myeloid cell subsets, including circulating monocytes, macrophages, and neutrophils(Lavin *et al*., 2014). KLF binding sites were enriched in genomic regions bound by the myeloid pioneering factor PU.1 in large cavity macrophages (LCM) (Gosselin *et al*., 2014), suggesting that KLFs may be involved in the development of these macrophages. Several groups have reported a role for KLFs in the activation and polarization of macrophages (Das *et al*., 2006; Das *et al*., 2012; Liao *et al*., 2011; Mahabeleshwar *et al*., 2011; Shi *et al*., 2014; Sweet *et al*., 2020; Roberts *et al*., 2017), but their contribution to early macrophage development and specification has not been characterized.

While resident macrophages exist in every tissue in the body, here we focus primarily on macrophage populations from the serous cavities and the lung. The appropriate induction of LCM in the peritoneal, pleural and pericardial serous cavities has previously been shown to require retinoic acid signaling and the expression of the transcription factor GATA6 (Buechler *et al*., 2019; Gautier *et al*., 2012; Gautier *et al*., 2014; Okabe and Medzhitov, 2014; Rosas *et al*., 2014; Gosselin *et al*., 2014; Casanova-Acebes *et al*., 2020). Development of alveolar macrophages (AM) in the lung is the result of early macrophages receiving GM-CSF (CSF-2) and TGF-β signals as well as the induction of the transcription factors C/EBPβ and PPARγ (Schneider *et al*., 2014; Dörr *et al*., 2022; Yu *et al*., 2017). Our group previously identified a role for two Group 2 KLFs, KLF2 and KLF4, in dampening innate immune responses against nucleic acids within apoptotic cell corpses when they are engulfed by LCM (Roberts *et al*., 2017). KLF2 was the dominant player in LCM; interestingly, KLF4 expression is correspondingly high in AMs(Mass *et al*., 2016), suggesting that KLF4 may be more functionally relevant in these cells. However, a broader role for these Group 2 KLF family members in the biology of these and other macrophage populations as well as their relationship to previously identified lineage-defining transcription factors remains unknown.

In this study we show that KLF2 and KLF4 are critically important for the development of LCM and AM, respectively. KLF2 deficiency results in a nearly complete absence of LCM *in vivo* due to an inability to recognize signals in the cavity environment, a phenotype distinct from what is observed in GATA6-deficient LCMs. We demonstrate that KLF2 directly regulates a number of LCM genes, including genes previously implicated in specifying LCM identity such as GATA6 and retinoic acid receptors. Similarly, deficiency in KLF4 impaired development of AM, and the remaining cells failed to express many known alveolar macrophage genes. Collectively, these results establish the importance of Group 2 KLFs for the induction of tissue-resident macrophage identities.

## RESULTS

### KLF2 is required for the development of serous Large Cavity Macrophages

The serous cavities contains two populations of macrophages commonly referred to as Small Cavity Macrophages (SCM; entirely monocyte derived) and LCM (embryonic derived, “tissue-resident” macrophages). These two populations are phenotypically distinct and easily identified by surface markers(Ghosn *et al*., 2010; Bain *et al*., 2016; Kim *et al*., 2016; Nguyen *et al*., 2012). SCM are CD11b^+^ F4/80^mid/lo^ MHCII^hi^ DNAM-1^hi^ ICAM2^-^ and LCM are CD11b^+^ F4/80^hi^ MHCII^-^ DNAM-1^-^ ICAM2^+^. Previous studies have shown a specific expression of GATA6 in LCMs relative to other tissue-resident macrophage populations (Lavin *et al*., 2014; Okabe and Medzhitov, 2014; Gosselin *et al*., 2014). Locally produced retinoic acid is required for GATA6 expression, and lack of GATA6 leads to a change in phenotype, function and the transcriptional landscape of LCM (Buechler *et al*., 2019; Okabe and Medzhitov, 2014; Casanova-Acebes *et al*., 2020). Based on the high expression of KLF2 in LCM, we sought to investigate the role of KLF2 relative to GATA6 in these cells. We analyzed immune cells isolated from the serous cavities of mice lacking KLF2 or GATA6 in myeloid cells by crossing *Klf2* floxed mice or *Gata6* floxed mice to LysMCre mice (*LysM*^Cre/+^ *Klf2*^fl/fl^, *LysM*^Cre/+^ *Gata6*^fl/fl^).

In unmanipulated adult wildtype B6 mice, most cavity macrophages (>80% of CD11b^+^F4/80^+^ cells) are LCM and less than 10% of macrophages are SCM (Bain and Jenkins, 2018). As previously described, GATA6 deficiency reduced the total percentage and number of macrophages (CD11b^+^F4/80^+^) in the serous cavities, which was mostly due to a reduction in LCM (Fig 1A, B). The remaining LCM expressed lower levels of F4/80, but otherwise expressed surface markers characteristic of wild-type LCM (Fig 1A). Thus, while reduced in number, the remaining *Gata6*-deficient progenitors did develop into LCMs, as previously described (Okabe and Medzhitov, 2014; Rosas *et al*., 2014; Gautier *et al*., 2014). *LysM*^Cre/+^ *Gata6*^fl/fl^ mice had a modest increase in frequency of SCM (Fig 1A,B), and the percentages and numbers of cavity monocytes (CD11b^+^Ly6C^+^) and neutrophils (CD11b^+^Ly6G^+^) were unchanged relative to wild-type controls (Fig S1A).

**Figure 1:**
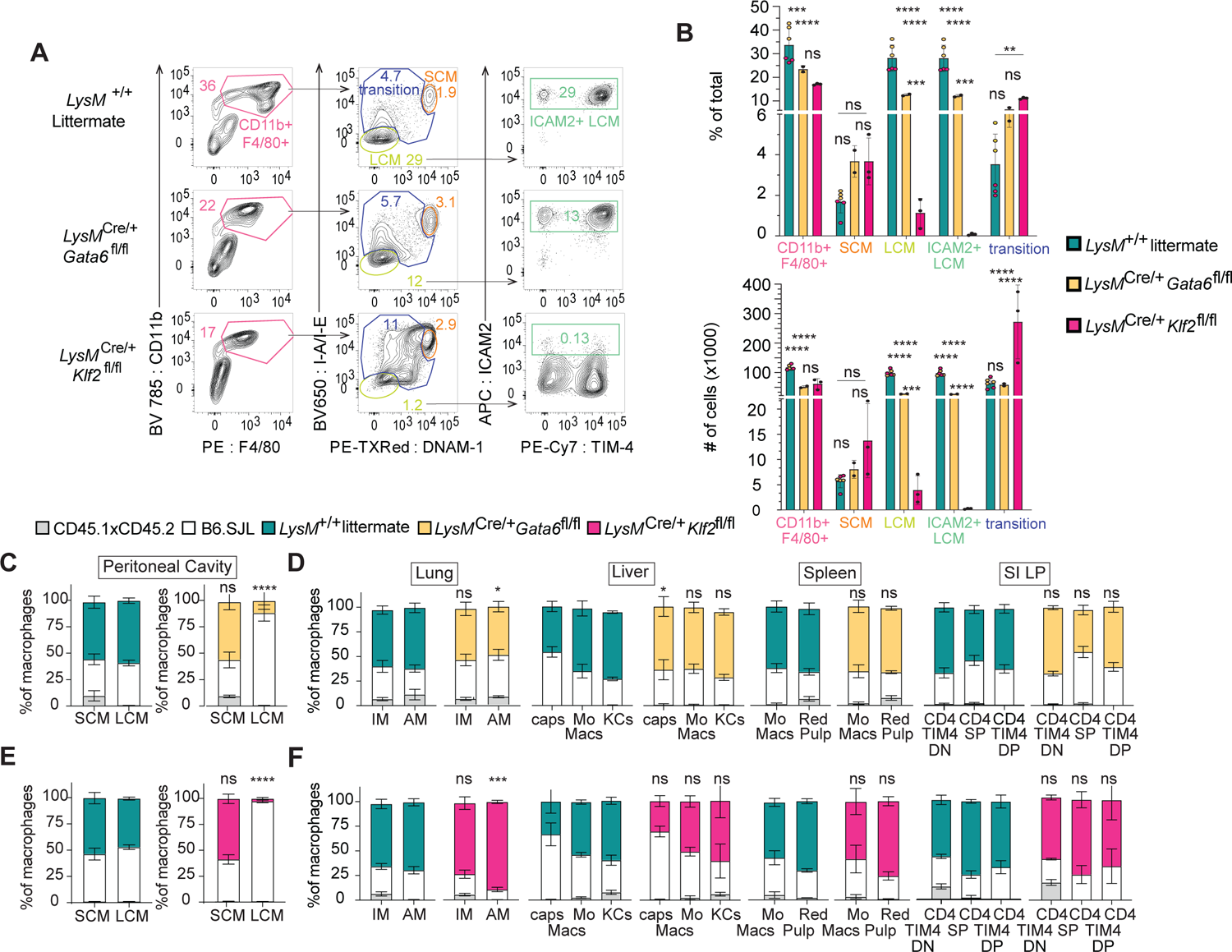
Impaired Development of Large Cavity Macrophages in LysMCre KLF2^fl/fl^ mice. **A:** Flow cytometry of cells in cavity lavage of *LysM^+/+^, LysM^Crel+^Klf2^¶m^*, and *LysM^Crel+^Gata6^nm^* mice, gated on live, single, CD3^−^B220^−^CD19^−^ cells. Numbers adjacent to gates indicate percent of total events. **B:** Bar graphs depict the percent of total and total number of CD11b^+^F4/80^+^, small cavity macrophages (SCM) (CD11b^+^F4/80^+^MHCIľDNAM-1^+^), large cavity macrophages (LCM) (CD11b^+^F4/80^+^MHCIĪDNAM-1^−^), and ICAM2* LCM, as measured by flow cytometry, **C:** Frequency of *LysM^+l+^Gata6^í^’^m^* or Lys/W^Creƒ+^Gaŕa6^flffl^ donor, B6.SJL donor, and CD45.1xCD45.2 host cells in SCM or LCM or **D:** monocyte and resident macrophage subsets in other tissues, **E:** Frequency of *LysM^+l+^Klf2^mt^* or *LysM^Crel+^Klf2^ĩm^* donor, B6.SJL donor, and CD45.1xCD45.2 host cells in SCM or LCM or **F:** monocyte and resident macrophage subsets in other tissues. Data are from one experiment representative of two independent experiments (A-F); for B, *LysM^+l+^* combined from both Gaŕa6^fl/fl^ and *Klf2^Wft^* controls, n=3 of each, LysM^Cre/+^GƏřa6^fl/fl^, n=2, *LysM^Cre/Ȼ^ Klf2^fl/fl^*, n=3; Significance determined by ordinary 2-way ANOVA with multiple comparisons and Šidák’s correction; C-F, mean and S.D. of three to four chimeras per group, Significance determined by Fisher’s exact test of relative contributions of each genotype in Bβ.SJL;Lys/W°^* chimeras compared to B6.SJL:Lys/W^+/+^ chimeras. Asterisks denote: ****<0.0001, *** 0.0006, ** 0.0021, * 0.033)

In contrast, *LysM*^Cre/+^ *Klf2*^fl/fl^ mice exhibited a severe reduction in LCM with a corresponding increase in SCM and a population of transitional cells with intermediate levels of DNAM-1 and MHCII (DNAM1^mid^ MHCII^mid^) (Fig 1A,B). The very few remaining CD11b^+^ F4/80^+^ MHCII^-^ DNAM-1^-^ cells did not express ICAM2 and had lower levels of F4/80, suggesting that they are not *bona fide* LCM and that no such cells exist in the serous cavities of *LysM*^Cre/+^ *Klf2*^fl/fl^ mice (Fig 1A, B). The frequency and absolute number of monocytes (CD11b^+^Ly6C^+^MHCII^+ or-^) was increased in *Klf2*-deficient mice and neutrophils were unchanged, showing KLF2 is not required for the development of either cell type (Fig S1A,B). We hypothesize that the increase in SCM, transitional cavity macrophages, and monocytes is due to the recruitment of cells to the cavity as a result of the “open” LCM niche (Fig S1B).

We also examined the requirement for KLF2 in LCM when competing with wildtype cells by making mixed bone marrow chimeras. Again, we included *Gata6*-deficient cells as a comparison. Both *Klf2*-deficient and *Gata6*-deficient cells failed to compete with wildtype cells in the generation of LCM, while SCM were equally comprised of wild-type and deficient cells (Fig 1C). The reduction of LCM was more complete for *Klf2*-deficient cells than *Gata6*-deficient cells (Fig 1C); however, the difference between the genotypes was minor when compared to the analysis of *LysM*^Cre/+^ *Klf2*^fl/fl^ and *LysM*^Cre/+^ *Gata6*^fl/fl^ mice at homeostasis (Fig 1A,B). In control mice that received a mixture of bone marrow from two wild-type mice, both the SCM and LCM populations had equal contributions from each donor. Importantly, loss of either GATA6 or KLF2 had no considerable effect on the development of any other tissue-resident macrophage populations in the liver, spleen, lung and small intestine. (Fig 1D, F, S1C-H). Altogether, these results establish an essential role for KLF2 in the development of LCMs that appears to be distinct from the role played by GATA6.

### KLF2 controls an LCM gene signature induced by the serous cavity environment

Tissue-resident macrophage populations from different tissues have distinct transcriptional profiles, and macrophages transferred into tissues can adopt the transcriptional state of that new environment (Lavin *et al*., 2014; Roberts *et al*., 2017). The inability of progenitors to differentiate into LCM in the absence of KLF2 suggests that KLF2 may control expression of specific genes required for LCM differentiation or maintenance in the serous cavities. However, the complete loss of LCM in *LysM*^Cre/+^ *Klf2*^fl/fl^ mice prevented any direct analysis of gene expression at homeostasis, so we chose to use a transfer model (Lavin *et al*., 2014; Roberts *et al*., 2017; Sakai *et al*., 2019) to examine KLF2-dependent gene regulation in macrophages transferred into the serous cavities (Fig 2A). *LysM*^+/+^ *Klf2*^fl/fl^ or *LysM*^Cre/+^ *Klf2*^fl/fl^ BMMs were transferred into congenically marked hosts whose cavities were pre-cleared by survival lavage to generate an open niche. After 3 or 7 days, transferred *LysM*^+/+^ *Klf2*^fl/fl^ BMMs upregulated ICAM2 by day 3 and further on day 7 (Fig 2B,C). In contrast, *LysM*^Cre/+^ *Klf2*^fl/fl^ BMMs upregulated MHCII but not ICAM2. Thus, the expression of KLF2 contributes to the ability of transferred macrophages to upregulate known phenotypic markers of LCMs, and in the absence of KLF2 they phenotypically resemble SCMs.

**Figure 2:**
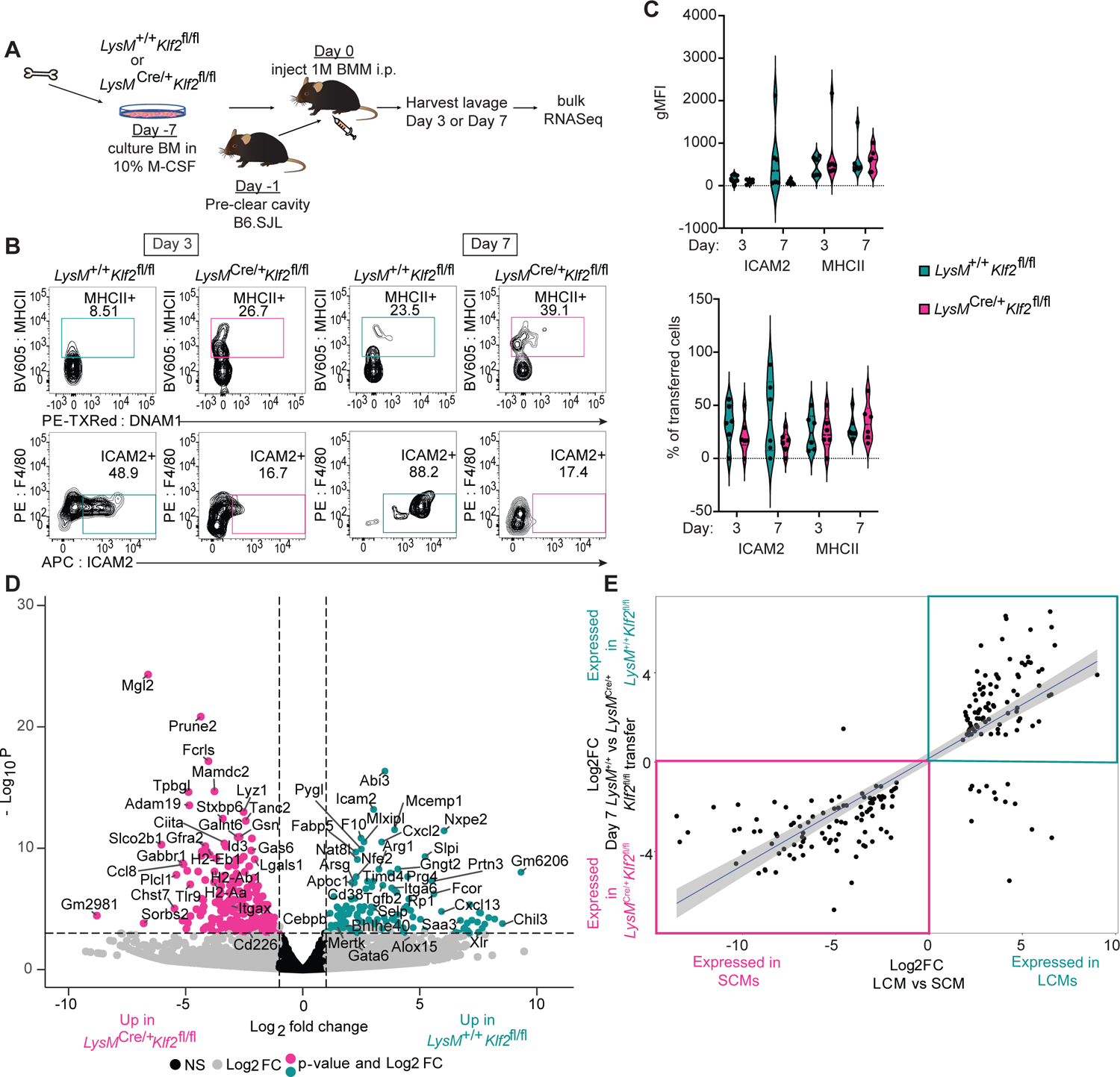
KLF2 is Required for Transferred Bone Marrow-Derived Macrophages to Rapidly Acquire Expression of Large Cavity Macrophage Identity **A:** Schematic of bone marrow macrophage (BMM) production, transfer into peritoneal cavity, harvest and analysis, **B:** flow cytometry of recovered and sorted *LysM^KIf^* or *LysM^Cre,+^Klf2^íìm^* BMMs on day 3 or day 7, gated on live, single, CD45.2^+^ cells. **C:** Violin plots depict the geometric MFI or the percent of transferred cells (CD45.2^+^) expressing the indicated cavity macrophage markers. **D:** Volcano plot of differentially expressed gene comparing sorted *LysM^+/+^Klf2^Wfí^* to *LysM^Cre/+^Klf2™* transferred BMMs. Genes with a Iog2 Fold Change greater than |11 are colored grey. Genes with a p-value of -Iog10 (p value) greater than 4 are above the horizontal black line. Genes meeting both criteria are colored: teal, enriched in *LysM^+/+^Klf2^m−^*, or pink, enriched in *LysM^Cre/+^Klf2^ím^.* **E:** Scatter plot of RNA-Seq expression data presenting the values of Iog2 Fold Change for genes differentially expressed in sorted small versus large cavity macrophages (X-axis) versus the genes differentially expressed between the day 7 sorted *LysM^+,+^Klf2^ím^* vs *LysM^Cre,+^Klf2^fí/fì^* transferred BMMs (Y-axis). Data are from one experiment representative of two independent experiments (B,C). Sequencing (D) was performed once. Cells shown in flow cytometry plots and violin graphs are from the same experiment that was sequenced. *LysM^+/+^Klf2^ím^*, n=5-6, *LysM^Cĩe/+^Klf2^ĩm^*, n=6.

We sorted *LysM*^+/+^ *Klf2*^fl/fl^ and *LysM*^Cre/+^ *Klf2*^fl/fl^ BMMs 7 days after transfer and performed bulk RNASeq to examine how lack of KLF2 impacted global gene expression in macrophages responding to tissue-derived signals. This analysis identified more than 594 differentially expressed genes (DEGs) on day 7 (fold change >2, false discovery rate<0.1) when comparing transferred *LysM*^+/+^ *Klf2*^fl/fl^ or *LysM*^Cre/+^ *Klf2*^fl/fl^ cells (Fig 2D). Transferred *LysM*^+/+^ *Klf2*^fl/fl^ BMMs expressed higher levels of many LCM identity genes including *Icam2*, *Timd4*, *Cebpb*, *Bhlhe40, Mertk*, *Cxcl13*, *Alox15*, and *Gata6*. *LysM*^Cre/+^ *Klf2*^fl/fl^ BMMs expressed relatively higher levels of genes previously identified in SCMs including MHCII genes (*H2-Aa*, -*Ab1*, *-Eb1*, *Ciita, etc*), *Itgax*, *Tlr9*, and *Cd226* (DNAM-1). We then compared these DEGs to those found in a dataset we generated from comparing LCM and SCM from wild-type mice. We found that the DEGs enriched in the recovered *LysM*^+/+^ *Klf2*^fl/fl^ BMMs correlated with genes with relatively greater expression in LCM compared to SCM (Fig 2E). Furthermore, like the endogenous *LysM*^Cre/+^ *Klf2*^fl/fl^ cavity macrophages, transferred *LysM*^Cre/+^ *Klf2*^fl/fl^ BMMs express genes that are enriched in SCMs (Fig 2E, SFig 2B). Importantly, only 38 genes, none of which were SCM or LCM signature genes, were differentially expressed (fold change >2, false discovery rate<0.1) between *LysM*^+/+^*Klf2*^fl/fl^ BMMs and *LysM*^Cre/+^*Klf2*^fl/fl^ BMMs before transfer into the cavity (SFig 2C), indicating that the DEGs detected in transferred BMMs were induced by the cavity environment. Taken together, these data support a model in which KLF2 activity is induced by cavity-dependent signals and is required for the expression of genes associated with LCM identity.

### Identification of KLF2 target genes involved in LCM identity

To identify which of the KLF2-dependent differentially expressed genes are directly regulated by KLF2, we performed CUT&RUN (Skene and Henikoff, 2017) on LCMs. To enhance the ability to pulldown KLF2-specific fragments we used an anti-GFP antibody on LCM from GFP-KLF2 transgenic mice(Weinreich *et al*., 2009). These mice have an N-terminal eGFP knocked into the endogenous KLF2 locus resulting in the expression of a fusion protein (SFig 3A). Additionally, the lower cell number requirement for CUT&RUN allowed us to generate biological replicates, further enhancing our ability to identify independently repeatable peaks with higher sensitivity as compared to ChIP-Seq. After filtering and trimming low quality bases, reads were mapped and peak analysis was performed with SEACR(Meers, Tenenbaum and Henikoff, 2019). Mapped reads were normalized to spike-in E. coli genome and we are reporting here the top 1% of those peaks.

**Figure 3:**
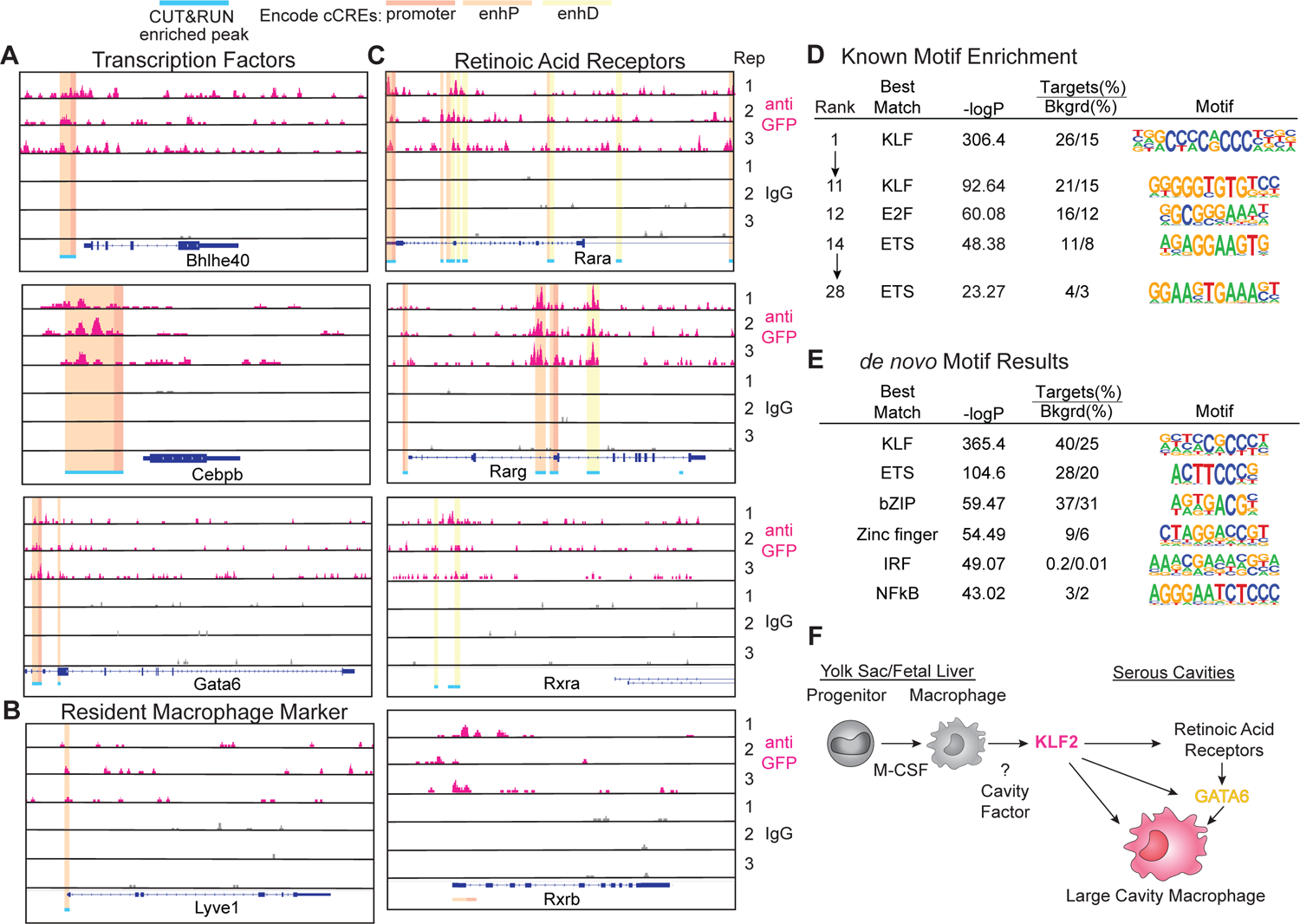
Lineage Determining Factors are Directly Regulated by KLF2 in Large Cavity Macrophages Genome browser tracks of anti-GFP KLF2 (pink) or IgG control (grey) CUT&RUN peaks from Large Cavity Macrophages at **A:** transcription factor, **B:** *Lyvel*, and **C:** retinoic acid receptor loci with top 1% of peak regions identified by SEACR in two out of three biological replicates marked in blue. ENCODE cCREs (candidate Cis Regulatory Elements) are highlighted to distinguish features occurring within the peak region (red: promoter, orange: proximal enhancer, yellow: distal enhancer). All genes are shown 5’->3’ and peak signals of 0-10 are shown on the y-axis. **D:** Known motif enrichment, and **E:** *de novo* motif enrichment analysis of Large Cavity Macrophage KLF2 CUT&RUN peaks compared to the IgG Control. **F:** Model of proposed function of KLF2 in Large Cavity Macrophages.

We found that KLF2 was bound to sites within or adjacent to several key LCM identity genes and nearly all of these loci overlapped with ENCODE-identified candidate cis-regulatory elements (cCREs; predominately proximal enhancer and promoter regions)(Moore *et al*., 2020). The genes bound by KLF2 included LCM transcription factors (*Bhlhe40*, *Cebpb* and *Gata6*) (Fig 3A) and surface markers like *Lyve1* (a marker that is important for transitioning to LCM (Han *et al*., 2024; Finlay *et al*., 2023)(Fig 3B), but we did not detect peaks in other LCM genes, including *Timd4, Icam2* or *Mafb (*SFig 3B). Additionally, the retinoic acid receptors (*Rara*, *Rarg*, *Rxra*) were enriched for KLF2 peaks (Fig 3C), confirming the importance of KLF2 in initiating the broader transcriptional identity of Large Cavity Macrophages.

Performing transcription factor motif analysis on the 1% of identified peaks revealed that the top 11 enriched known motifs all matched to KLF binding sites, confirming the validity of the data with additional representation of E2F binding sites and ETS transcription factor motifs (Fig 3D). When we performed *de novo* motif discovery on total enriched peaks, KLF binding sites were the most enriched followed by ETS, with lesser contribution from sites for bZIP, Zinc finger, IRF and NFκB transcription factors (Fig 3E). Critically, genes directly bound by KLF2 were not co-enriched for GATA6 or RAR/RXR binding motifs, consistent with our model that KLF2 is required for the responsiveness of developing LCM to retinoic acid (Fig 3F). These analyses demonstrate the importance of KLF2 for direct regulation of expression of known LCM transcription factors (i.e., GATA6 and RARs) while also potentially collaborating with general macrophage transcription factors like PU.1 (ETS) and AP-1/ATF/CEBP (bZIP). The paramount role of KLF2 in the transcriptional network controlling LCM identity (Fig 3F) likely explains the more severe effect on LCM development observed with KLF2 deficiency relative to Bhlhe40 deficiency(Jarjour *et al*., 2019; Rauschmeier *et al*., 2019), C/EBPβ deficiency(Dörr *et al*., 2022; Cain *et al*., 2013) or GATA6 deficiency (reduction or altered phenotypes of LCMs).

### Ectopic expression of KLF2 induces LCM identity *in vitro*

Our previous experiments identified genes that are differentially expressed between SCMs and LCMs but could not distinguish between transcripts regulated by transcription factors from those that are regulated by cavity-dependent signals. To identify specific genes whose expression is induced by the expression of KLF2, we employed a simplified model of overexpression (OE) by retroviral (RV) transduction in *in vitro*-derived BMMs (Fig 4A).

**Figure 4:**
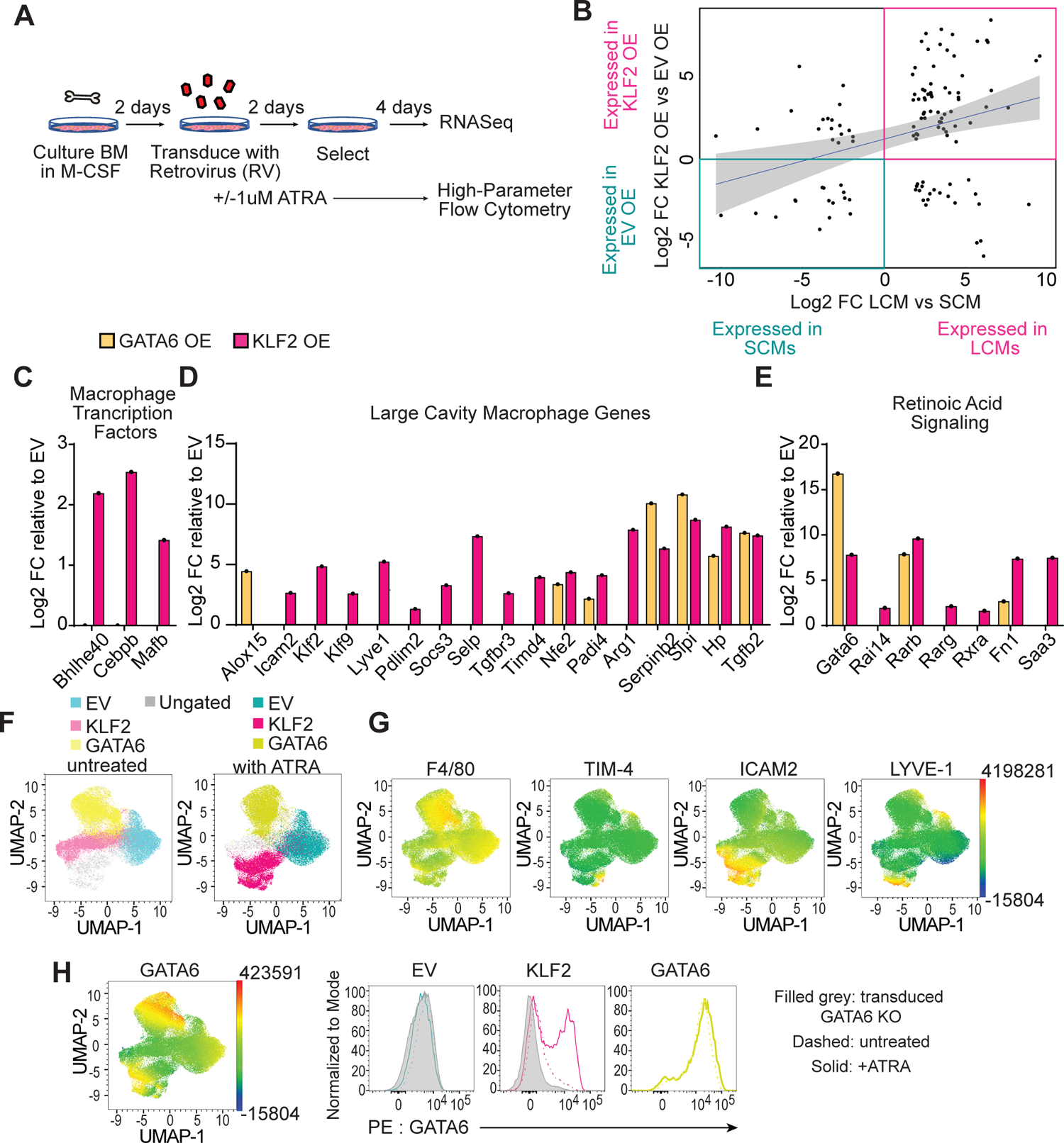
Ectopic Expression of KLF2 Confers Large Cavity Macrophage Phenotype and Transcriptional Identity *In Vitro*. **A:** graphical representation of retroviral (RV) overexpression (OE) of transcription factors in in vitro-derived bone marrow macrophages. **B:** Scatter plot of RNA-Seq expression data presenting the values of Iog2 Fold Change for genes differentially expressed in sorted large versus small cavity macrophages (X-axis; same as Fig 2D) versus the genes differentially expressed between the EV versus KLF2 transduced BMMs (Y-axis), **C-E:** Bar graphs representing Iog2 Fold Change of RNA-Seq data from transcription factor transduced BMMs relative to EV. **C:** large cavity macrophage relevant transcription factors, **D:** other relevant large cavity macrophage genes, and **E:** retinoic acid receptors and response genes. **F:** UMAP plot of GFP (virus)+ cells. Live, single, virus+ cells from empty vector (EV), KLF2 OE or GATA6 OE were gated, downsampled to 12879 cells per sample which were barcoded and concatenated. Live/dead stain and GFP were excluded from the list of UMAP running parameters. The resulting UMAP projection is colored according to overlaid gating based on transduction and treatment. **G:** UMAP plots showing expression intensities of large cavity macrophage markers. **H:** UMAP plot showing expression intensity of GATA6 and same data shown as histograms. Filled grey is RV transduced *LysM^Crel+^Gata6™* BMMs, dashed line is RV OE, and solid line is RV OE plus ATRA. Dashed and solid lines are transduced B6 BMM. (Data are from one experiment representative of two independent experiments (F,G,H). RNA-Seq was performed on samples generated in three independently performed transduction experiments. RNA was harvested, libraries were generated, and sequencing was performed concurrently.

We first analyzed what transcripts were induced by the expression of KLF2 or GATA6 relative to the empty vector (EV) control by bulk RNASeq. Similar to our analysis of transferred BMMs, we compared the DEGs from the comparison of KLF2 OE to the EV control with DEGs between LCM and SCM. Strikingly, overexpression of KLF2 alone, without any additional tissue-specific soluble factors or any cavity-resident cells, can recapitulate the induction of many LCM-specific genes (Fig 4B - E). Notably, overexpression of KLF2, but not GATA6, induced the expression of LCM transcription factors including *Bhlhe40*, *Mafb*, and *Cebpb* (Fig 4C). KLF2 also induced many canonical LCM identity genes. A subset of these genes were induced only by KLF2 (e.g., *Icam2*, *Lyve1*, *Pdlim2*, *Socs3*) (Fig 4D), while others were induced by both KLF2 and GATA6 (e.g., *Serpinb2*, *Hp*, *Tgfb2*). The difference of expression between these two groups of genes is most likely due to the fact that KLF2 also induced the expression of retinoic acid receptors (*Rarb*, *Rarg*, *Rxra*) and *Gata6* (Fig 4E), which can in turn induce expression of retinoic acid-dependent LCM identity genes (e.g., *Serpinb2*, *Hp*, *Tgfb2)*. Thus, KLF2 expression is sufficient to induce the expression of many LCM identity genes, including the retinoic acid/GATA6 axis previously identified as critical for the specification and function of these cells (Fig 3G).

To further test the hypothesis that KLF2 OE induces a macrophage state that more closely recapitulates LCM than GATA6 OE, we used high-dimensional flow cytometry with an 18-parameter myeloid cell panel. Transduced cells were cultured in the presence or absence of exogenously added all-trans retinoic acid (ATRA) to confirm the functional importance of induced retinoic acid receptor expression and to determine if exogenous ATRA could enhance adoption of an LCM-like phenotype. When we concatenated equal events from all conditions and performed UMAP analysis, the three transduced BMM populations clustered separately from each other, indicating that overexpression of KLF2 or GATA6 distinctly altered the phenotype of BMMs. (Fig 4F) The addition of ATRA changed both the empty vector and the KLF2 OE transductants but had little effect on the GATA6 OE cells (Fig 4F), consistent with GATA6 being downstream of retinoic acid signaling (Okabe and Medzhitov, 2014; Buechler *et al*., 2019). When we examined known LCM marker expression, KLF2 OE cells were the predominate population that gained expression of markers (ICAM2, LYVE-1, and TIM-4) that phenotypically define LCMs or transitional LCMs (Han *et al*., 2024; Finlay *et al*., 2023). The expression of these markers was further enhanced by ATRA only in KLF2 OE BMMs (Fig 4G), suggesting that KLF2 expression precedes LCM identity generation, including the ability to receive retinoic acid signals (Fig 3G). Detection of GATA6 protein by intracellular staining confirmed induction in KLF2 OE cells which was further enhanced by addition of ATRA (Fig 4H). Together the RNASeq and high-parameter flow cytometry show that the expression of KLF2 and the addition of ATRA synergize to induce and enhance expression of LCM markers *in vitro*.

### Appropriate induction of alveolar macrophage identity requires KLF4

Next, we considered whether the requirement for KLF2 in LCM reflects a more general function of KLFs in regulating the development and identity of resident macrophage populations in other tissues. KLF4 is preferentially expressed in AM (Mass *et al*., 2016), so we examined whether deficiency in KLF4 would affect their development. Conditional knockout (KO) of KLF4 with either *LysM*- or CD11c-Cre led to a reduction in frequency and number of AM, while leaving other myeloid populations relatively unaffected (Fig 5A, B). Moreover, in mixed bone marrow chimeras, *LysM*^Cre/+^ *Klf4*^fl/fl^ cells were outcompeted by B6.SJL wild-type cells in AMs while both genotypes were equally represented among the monocyte-derived interstitial macrophage population of the lung (Fig 5D), similar to what has been reported for PPARγ-deficient or C/EBPβ-deficient+wild-type mixed bone marrow chimeras (Schneider *et al*., 2014; Dörr *et al*., 2022). KLF4 deficiency did not have any major impact on macrophage subsets in spleen, liver or small intestine.

**Figure 5:**
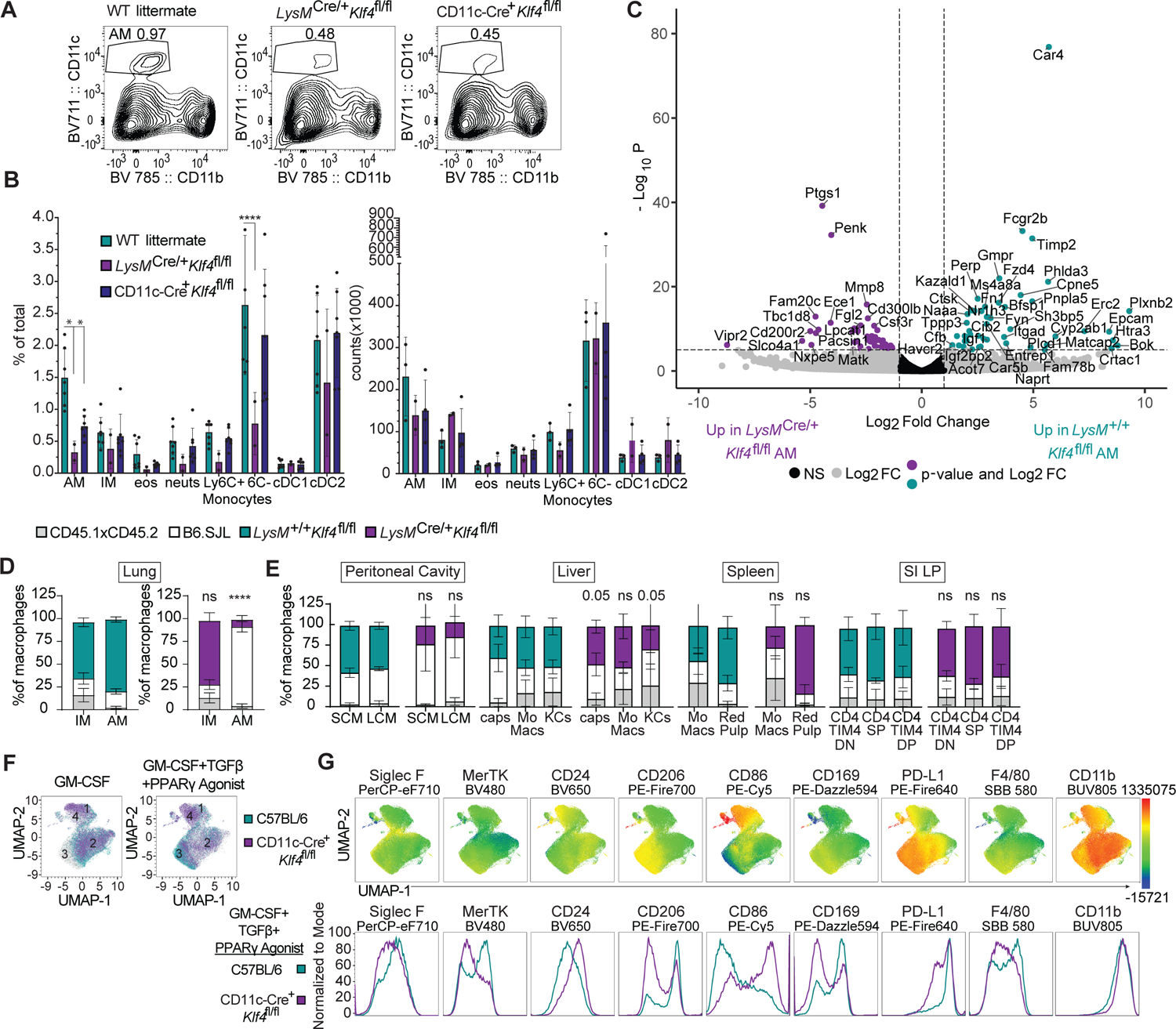
KLF4 is Important for Alveolar Macrophage Development and Identity A: Flow cytometry of cells in lungs of wild-type littermate, *LysM^Cre+^Klf4^fl/fl^*, and CD11c-Cre+/*Klf*4^fl/fl^ mice, gated on live, single, CD45^+^CD3^−^B220^−^CD19^−^ cells. Numbers adjacent to gates indicate percent of total events. B: Bar graphs depict the percent of total and total number of alveolar macrophages (AM; CD11c^+^CD11b^−^, Siglec-F^+^, CD24), interstitial macrophages (IM: CD11b^+^, MHClľ, CD24’, CD64^+^), eosinophils (eos; MHClľ, CD11b^+^, CD24^+^, Siglec-F^+^), neutrophils (neuts; MHCIĪ, CD11b^+^, CD24^+^, Ly6G^+^), Ly6C^+^ and LyθC^−^ monocytes (MHCIĪ, CD11b^+^, CD64^+^, Ly6C^+^ or Ly6C^mid/−^), cDC1 (CD11c^+^,CD11b-, Siglec-F^−^, CD24^+^) and cDC2 (CD11b^+^, MHCII^+^, CD24^+^, CD64^−^) dendritic cells (DCs) as measured by flow cytometry (Misharin, 2013), C: Volcano plot of differentially expressed gene comparing sorted *LysM^+l+^Klf4^ĨWI^* to *LysM^Crel^*Klf4^wt^’* alveolar macrophages. Genes with a Iog2 Fold Change greater than |11 are colored grey. Genes with a p-value of -Iog10 (p value) greater than 6 are above the horizontal black line. Genes meeting both criteria are colored: teal, enriched in *LysM^KIf^;* purple, enriched in *LysM^Crel+^Klf4^w¶^).* D: Frequency of *LysM^+l+^Klf4^m^’* or *LysM^Crel+^Klf4^fWİ^* donor, B6.SJL donor, and CD45.1xCD45.2 host cells in alveolar and interstitial macrophages in the lung or E: monocyte and resident macrophage subsets in other tissues. F: UMAP plot of live, single cells were gated, downsampled to 22758 cells per sample which were barcoded and concatenated. Live/dead was excluded from the list of UMAP running parameters. The resulting UMAP projection was overlayed with gates representing the genotype (C57BL/6 or CD11c-Cre+K/f4^fl/fl^) and culture conditions (GM-CSF alone or GM-CSF+TGFb+PPARg agonist). G: UMAP plots showing expression intensities of various macrophage and dendritic cell markers (top row) and those same markers shown as histograms from UMAP data of just the GM-CSF+TGFb+PParg agonist culture condition (bottom row). Data are combined from two independent experiments (A, B, D, E), significance determined by ordinary 2-way ANOVA with multiple comparisons and Šidák’s correction; Bulk RNA Sequencing (C) was performed from one set of samples containing two biological replicates of each genotype; (D,E), mean and S.D. of three to four chimeras per group from one representative of two independent experiments, significance determined by Fisher’s exact test of relative contributions of each genotype in B6.SJL:/_ys/ư^Cre+^K/f4^fl/fl^ chimeras compared to *BQ.SĴL·LysM^+l+^Klf4^fim^* chimeras. Asterisks denote: ****<0.0001, *** 0.0006, ** 0.0021, * 0.033)

We sorted AM from *LysM*^+/+^ *Klf4*^flfl^ and *LysM*^Cre/+^ *Klf4*^fl/fl^ mice for RNA-Seq. Several known alveolar macrophage identity genes were found to be enriched in wild-type cells, the most differentially expressed included *Car4*, *Perp*, *Epcam*, *Cpne5*, *Fcgr2b* (CD64), *Cd206*, *Cd74*, and *Ctsk* (Fig5 C). GO enrichment analysis of the genes preferentially expressed in the wild-type AMs resulted in terms related to cell proliferation, lipid, fatty acid and carboxylic acid metabolism, and antigen presentation via MHCII (SFig 3C). These pathways and processes have previously been recognized as important to AM function and were also enriched in the differential gene signatures comparing wild-type AMs to AMs lacking PPARγ or C/EBPβ(Dörr *et al*., 2022). However, MHCII antigen presentation related genes are enriched in C/EBPβ-deficient AMs, not wild-type, suggesting that C/EBPβ and KLF4 have distinct functions in AMs. The *LysM*^Cre/+^ *Klf4*^fl/fl^ AMs have higher relative expression of genes related to cell adhesion and mobility as well as genes involved in positive regulation of the immune system and cytokine signaling (SFig 3D). The expression of these genes in *LysM^C^*^re/+^ *Klf4*^fl/fl^ AMs may indicate that these altered AMs represent newly arriving cells that are trafficking to the alveolar space due to the partially unfilled niche, similar to the increase in SCM and transitional macrophages that we observe in the serous cavities in *LysM*^Cre/+^ *Klf2*^fl/fl^ mice.

Based on these *in vivo* results, we asked whether KLF4 deficiency would prevent the development of *in vitro*-derived AM-like cells by culturing bone marrow progenitors from CD11c-Cre+ *Klf4*^fl/fl^ or wild-type C57BL/6 mice in factors that induce AM development and KLF4 expression: GM-CSF, TGF-β and the PPARγ agonist rosaglitizone (Gorki *et al*., 2022; Luo *et al*., 2021). After nine days, we used high-parameter flow cytometry to evaluate the impact on macrophage differentiation. In the presence of GM-CSF alone, most wild-type and CD11c-Cre+ *Klf4*^fl/fl^ cells clustered together (Cluster 2), though a population of cells cluster separately (Cluster 1) and contains a majority of CD11c-Cre+ *Klf4*^fl/fl^ cells. When TGF-β and PPARγ agonist were added, most wild-type cells differentiated into a distinct cluster of cells (Cluster 3). The majority of CD11c-Cre+ *Klf4*^fl/fl^ cells did not convert fully to Cluster 3, stopping just short and many remaining in Cluster 4 (adjacent to Cluster 1). (Fig 5F). In wild-type cells, GM-CSF+TGF-β+PPARγ agonist induced upregulation of alveolar macrophage markers (Siglec-F, MerTK, CD206, CD169, F4/80), while suppressing “dendritic cell”/non-AM markers (CD86, CD11b). In the absence of KLF4, AM markers were induced to a lesser extent, and “dendritic cell” markers were maintained (Fig 5G), suggesting that KLF4 is necessary for progenitor cells to develop into alveolar macrophages in response to known required AM soluble signals, similar to what we observed with KLF2 *in vitro.* These results support a model in which KLF4 is required for the development of AMs.

## DISCUSSION

Tissue-resident macrophages perform important functions that maintain homeostasis in their tissue of residence. These macrophages develop during embryogenesis and are instructed through local signals to gain tissue specific identities. Over the past decade, extensive work from many labs has begun to identify the key transcription factors and molecular pathways that are important for each resident macrophage population, but how such factors and pathways cooperate to determine a given macrophage identity remains unclear and is an area of intense investigation. In this study, we establish that members of the Group 2 Krüppel-like Factor family are critical for development of two specific tissue-resident macrophage populations.

Previous experiments have identified retinoic acid receptor signaling and GATA6 as essential transcriptional regulators of embryonic-derived LCM (Buechler *et al*., 2019; Finlay *et al*., 2023; Gautier *et al*., 2012; Gautier *et al*., 2014; Okabe and Medzhitov, 2014; Rosas *et al*., 2014; Casanova-Acebes *et al*., 2020). Here we show, in a side-by-side comparison, that KLF2 deficiency has a more dramatic effect on the development of LCM than GATA6 deficiency. in a cell-intrinsic, tissue-specific manner. Through multiple lines of evidence, we determine that the induction of KLF2 is required when tissue-residency is established and not during earlier stages of macrophage development. We show that KLF2 promotes the induction of other transcription factors important in LCM, including GATA6, and enables increased capacity for retinoic acid sensing through direct regulation of retinoic acid receptors, which also control GATA6 expression. Together these data place KLF2 upstream of previously identified regulators of LCM development and demonstrate that KLF2 is sufficient to induce many of the critical genes required for LCM development.

KLF2 is necessary for retinoic acid-dependent elements of LCM identity but is also essential for many of the LCM genes that are retinoic-acid independent, by both direct and indirect mechanisms (not all genes we observed as being transcriptionally regulated were also bound directly by KLF2 in our CUT&RUN). For example, reduced TLR signaling is a distinct module of LCM identity that is KLF2-dependent and retinoic acid-independent (Roberts *et al*., 2017). Though retinoic acid signaling and GATA6 are important for several identified LCM functions (Okabe and Medzhitov, 2014; Finlay *et al*., 2023; Casanova-Acebes *et al*., 2020; Deniset *et al*., 2019), *Gata6*- or retinoic acid receptor-deficient LCMs express KLF2, and thereby, are present in greater numbers than in *Klf2*-deficient mice, and have a fairly normal LCM phenotype.

While the cells responsible for retinoic acid production in the cavity have been identified, it remains unknown how KLF2 expression is regulated in LCMs. *Klf2* expression is induced by laminar flow in endothelial cells or by blocking HMG-CoA reductase in endothelial and T cells via the induction of the transcription factor MEF2 (Maejima *et al*., 2014; Parmar *et al*., 2006; Villarreal *et al*., 2010; Ahmad and Lingrel, 2005). It is possible that macrophage progenitor cells in the cavity experience a specific flow rate or that the cavity induces a change in metabolism that activates MEF2, either through HMG-CoA accumulation or inhibition of cholesterol synthesis. The relevance for these potential mechanisms in LCM will be important to address in future work. Of course, distinct undiscovered mechanisms may also be involved in KLF2 regulation in these cells.

Our work also establishes the importance of KLF4, a different Group 2 KLF, in AM development. While the precise mechanisms of KLF4 function in these cells require further investigation, our results demonstrate that expression of many AM genes is altered in the absence of KLF4. Previous studies have demonstrated the importance of GM-CSF and TGF-β signaling to induce C/EBPβ and PPARγ expression in AM (Dörr *et al*., 2022; Schneider *et al*., 2014; Yu *et al*., 2017). What regulates KLF4 in these cells and whether KLF4 regulates these previously identified pathways or other aspects of AM development remains unclear. It is intriguing that both subsets of macrophages we have identified as having Group 2 KLF-dependency also both depend on C/EBPβ. Because of this concordance, we speculate that KLF4 may also act upstream of C/EBPβ in AMs as KLF2 does in LCM.

The requirement of individual KLF family members for development of specific tissue-resident macrophage populations is somewhat unexpected, especially considering their rather broad expression across many cell types. However, we contend that this characteristic may explain why their role in macrophage function was previously missed. When looking for transcriptional regulators that are critical for the identity of a particular macrophage population, investigators have focused on genes with narrow, population-specific expression. This approach may miss genes with broader expression that nevertheless play critical regulatory functions in particular cells. Another approach to identify transcription factors important for macrophage development is motif analysis of histone marker ChIP- or ATAC-Seq datasets. Through these types of experiments, KLF binding sites have been noted to be enriched at genomic loci of active genes of various tissue-resident macrophages. However, KLF binding sites were also enriched in the total enhancer regions of neutrophils and monocytes, further supporting the idea that KLF regulation is a common feature of myeloid cells. Finally, this kind of computational analysis of open chromatin or enhancer regions can only identify KLF binding motifs in the broadest sense (including in our own data, Fig 3D), as all KLFs share the same core binding motif and specific binding site sequences for individual KLF family members have not been determined.

Finally, our data ascribe a new role for KLFs in macrophages that is different than what has been previously reported. Previous work has suggested that KLF2 and/or KLF4 limit the activation and proinflammatory functions of monocytes and macrophages within various tissue and in various disease models (Chakraborty *et al*., 2023; Liao *et al*., 2011; Roberts *et al*., 2017; Das *et al*., 2006; Das *et al*., 2012; Mahabeleshwar *et al*., 2011; Shi *et al*., 2014; Sweet *et al*., 2020; Herta *et al*., 2021; Manoharan *et al*., 2019). While our analyses do not directly refute an anti-inflammatory role for KLF2 and KLF4, we do believe that such effects on activation may be secondary to the role that these transcription factors play in macrophage differentiation within tissues. For example, the inability of infiltrating KLF2-deficient monocytes to respond to tissue-derived cues in the inflamed serous cavities may result in activated monocyte-derived macrophages, but this gene signature may reflect the absence of KLF2-driven differentiation more than a direct regulation of macrophage activation.

In summary, we show that commonly expressed transcription factors belonging to the Group 2 KLF family are required for the development of distinct tissue-resident macrophage populations. In LCM, this requirement is based on KLF2-dependent expression of developmental regulators that enable cells to sense tissue-derived signals. Our results suggest that future work should consider potential roles for more broadly expressed transcription factors in development and identity of resident macrophages.

## MATERIALS AND METHODS

### Mice

All animal procedures were approved by the University of California, Berkeley Institutional Animal Care and Use Committee in accordance with University of California, Berkeley research guidelines for the care and use of laboratory animals. The following mice were used in this study: C57BL/6J (000664), B6.SJL (B6.Ptprc^a^Pep^c^/BoyJ, 002014), *LysM*^Cre^ (Lyz2Cretm1(Cre)Ifo, 004781) *Klf2*^fl/fl^ (gift from Jerry B Lingrel), *LysM*^Cre^ *Gata6*^fl/fl^ (gift from Paul Kubes, UCalgary), *LysM*^Cre^ *Klf4*^fl/fl^ (floxed *Klf4* allele: MMRRC 29877), CD11c-Cre *Klf4*^flfl^ (Tg(Itgax-cre)1-1Reiz/J, 008068, cross generated in house), GFP-KLF2 (gift from Stephen Jameson and Kristin Hogquist labs, UMinnesota), CD45.1 x CD45.2 F1 (generated in house by crossing C57BL/6J to B6.SJL (B6.Ptprc)). Mice were used between 7-9 weeks of age, except chimeric mice which were used 8-12 weeks after bone marrow injection.

### Cell preparation for flow cytometry

Peritoneal lavage was obtained by injecting 5mL PBS (Gibco) containing 2mM EDTA (Thermo) and 2% FCS (Gibco or Omega) with a 23 gauge needle into the peritoneal cavity and then retrieving the solution back out with the same syringe and a 21 gauge needle. Pleural lavage is performed the same, but with 3mL PBS containing 2mM EDTA and 2% FCS.

The spleen, thymus, kidney, lung and liver were harvested and homogenized into a single-cell suspension with C tubes on a GentleMacs (Milltenyi) and correct tissue programs. For spleen, thymus and kidney, LiberaseTM (Roche) and DNaseI (Sigma) were used in 2mL HBSS (Gibco) with Ca+ Mg+ and the same concentration was used for liver in 5mL HBSS with Ca+ Mg+. For lung, LiberaseTM and DNaseI in HBSS with Ca+ Mg+ in 2mL. Tissues were incubated 20min at 37 deg C and then finished on the GentleMacs. Enzymes were quenched with PBS with 2mM EDTA and 2% FCS on ice and then red blood cells were lysed with room-temperature ACK lysis buffer (Gibco) and passed through a 100μm cell strainer.

For intestinal LP cells preparation (Bergsbaken and Bevan, 2015), Peyer’s patches were removed with scissors, and the intestine was cut open longitudinally. The intestinal contents and mucus were removed by gentle scraping with forceps and rinsing with PBS. The intestine was cut into 1cm pieces and incubated in HBSS buffer containing 1mM dithiothreitol (Fisher) and 10% FCS at 37°C while stirring for 20min to remove IELs, which were discarded. Intestinal tissue was transferred to HBSS (Gibco) containing 1.3mM EDTA (Thermo) and stirred at 37°C for 20mins to remove the epithelium, which was discarded. Remaining intestinal pieces were then incubated in HBSS containing 5% FBS and 150U/ml collagenase type 2 (Worthington) at 37°C with stirring for 45min to isolate LP cells. LP cells were further purified by gradient centrifugation at 2800 rpm for 20min with 44% and 67% Percoll (GE Healthcare/Cytiva) at room-temperature. Afterwards, the mucoid top layer of dead cells and epithelial cells is removed. All the remaining Percoll, except the dead cell and red blood cell pellet, was collected and washed, as intestinal macrophages do not accumulate within the leukocyte layer at the interface.

### Flow cytometry

Dead cells were stained using LIVEDEAD (Thermo) for 15 min on ice in PBS. For traditional flow cytometry, nonspecific antibody binding was blocked by incubation with an anti-CD16/32 antibody (clone 2.4G2; UCSF Ab production facility) and whole mouse IgG (Sigma) on ice for 15 min, and the cells were stained with fluorophore-conjugated antibodies on ice for 20 min. Cells were kept on ice and analyzed on a BD Fortessa (BD Biosciences). For spectral flow, cells were incubated with anti-CD16/32 APC-Fire750 and anti-CD64 PECy7 for 15 min on ice before the addition of the rest of the fluorophore-conjugated antibodies were added on ice for 20 min in Brilliant Stain Buffer (BD Biosciences). For GATA6 staining, cells were fixed and permeabilized with the eBioscience™ Foxp3 / Transcription Factor Staining Buffer Set (Thermo) for 30 minutes followed by staining with GATA6-PE (Cell Signaling Technologies) at 1:400 for 30 minutes on ice. Cells were kept on ice until analyzed on a 5 laser Cytek Aurora (Cytek Biosciences). Data were analyzed in FlowJo (FlowJo, LLC). For UMAP analysis, equivalent numbers of cells in each were concatenated and all markers were included except LiveDead and GFP (for the retroviral transductants).

### Radiation bone marrow chimera generation

Recipient CD45.1 x CD45.2 F1 mice were lethally irradiated with a split dose of 500 rads in the evening of Day-1 and 450 rads on the morning of Day 0 using an XRad 300 X-ray producing machine (Precision Xray). Femur, tibia, and iliac bones from *LysM*^Cre/+^ *Klf2*^fl/fl^, *LysM*^+/+^ *Klf2*^fl/fl^ *LysM*^Cre/+^ *Gata6*^fl/fl^, *LysM*^+/+^ *Gata6*^fl/fl^, *LysM*^Cre/+^ *Klf4*^fl/fl^, CD11c-Cre+ *Klf4*^fl/fl^, *Klf4*^fl/fl^ and B6.SJL mice were harvested, flushed with RPMI and red blood cells were lysed with ACK buffer. Cells were counted and mixed 1:1 in appropriate combinations for the experiment. Cells were injected i.v. into the tail vein of recipient mice in 100uL of RPMI (Gibco). Mice were given 6 or 8 million total cells and kept cleanly without antibiotics in the presence of pre-irradiation fecal pellets for 8-12 weeks.

### Bone Marrow Macrophage culture and transduction

On Day-1, 2.5 million 293T-GP2 retroviral producer cells were plated in 10mL DMEM with HEPES (Gibco), Sodium Pyruvate (Gibco), Glutamax (Gibco), Pen/Strep (Gibco), B-mercaptoehtanol(Fisher), 10% FCS (Gibco or Omega)(DM10+) in 10cm TC treated plates. On Day 0, 293T-GP2s (Clontech/Takara, RRID:CVCL_WI48) were transfected with 100ng MSCV plasmid (MSCV multiple cloning site IRES puroR 2A mCherry or GFP) and 10ng pVSVg plasmid (Addgene 8454) with Lipofectamine 2000 (Thermo). Also on Day 0, femur, tibia and iliac bones from C57BL/6 mice were flushed with RPMI (Gibco) and red blood cells were lysed with ACK buffer (Gibco). Cells were counted and 4 million cells were plated in 10 cm non-TC treated plates in 8 mL RPMI with HEPES (Gibco), Sodium Pyruvate (Gibco), Glutamax (Gibco), Penicillin/Streptomycin (Gibco), B-mercaptoethanol (Fisher), 10% FCS (Gibco or Omega) (RP10+), and 10% titrated M-CSF 3T3-cell conditioned laboratory-made media. On Day 1, media was replaced on the 293T-GP2 to 8 mL. On Day 2 viral supes were harvested from 293T-GP2s and filtered through 0.45uM PES. Bone marrow cultures were harvested, resuspended in 293T-GP2 supernatant supplemented 10% M-CSF 3T3 cell media and returned to the same 10 cm plate. Media was replaced on 293T-GP2s for repeating of viral infections on Day 3. On Day 4, bone marrow cells are harvested out of plates and resuspended in 8 mL RP10+ with 10% M-CSF 3T3 supe with 5 ug/mL puromycin. Media was again replaced on Day 7 with fresh RP10+ with 10% M-CSF 3T3 supe. For concurrent treatment with all-trans retinoic acid (ATRA, Sigma), cells were given 1uM ATRA daily starting on Day 3. Cells were harvested on day 8-10 by incubating in 5mL 4mM ETDA in PBS for 20 min in a 37 deg C incubator. ETDA was quenched by the addition of 7mL RP10+. Cells were counted and 0.1 million cells were resuspended in TRIzol (Thermo) and stored at −80 for RNA isolation. The rest of the cells were analyzed by flow cytometry.

### Bone Marrow Macrophage culture and transfer

On Day-7, femur, tibia and iliac bones from *LysM*^Cre/+^ *Klf2*^fl/fl^, *LysM*^+/+^ *Klf2*^fl/fl^ mice were flushed with RPMI (Gibco) and red blood cells were lysed with ACK buffer (Gibco). Cells were counted and 1M cells were plated in 10cm non-TC treated plates in 8 mL RPMI with HEPES (Gibco), Sodium Pyruvate (Gibco), Glutamax (Gibco), Pen/Strep (Gibco), B-me(Fisher), 10% FCS (Gibco or Omega) (RP10+), 10% titrated M-CSF 3T3-cell conditioned laboratory-made media. On Day-3, BMMs were given an additional 8mL RP10+ with 10% M-CSF. On Day-1, B6.SJL recipient mice were anesthetized with (mg/kg) ketamine i.p. Upon achieving the appropriate anesthetic level, 5mL of warm PBS was injected slowly into the peritoneal cavity with a 23 gauge needle. Mice were gently rocked and then PBS was recovered with a 21 gauge needle. Mice were kept warm during the duration of the anesthesia and monitored until recovered. On Day 0, BMMs were harvested by incubating in 5mL 4mM ETDA in PBS for 20 min in a 37 deg C incubator. ETDA was quenched by the addition of 7mL RP10+. Cells were counted and resuspended to inject 1M cells per recipient in 100uL of RPMI i.p. Cells were harvested by lavage on day 3 or 5, stained and sorted on live, single CD45+ cells, with additional collection of data on expression of F4/80, CD11b, ICAM2, MHCII, TIM-4 and DNAM-1 (did I dump Bs?) with a BD FACS Aria Fusion (BD Biosciences). TRIzol LS (Thermo) was added to sorted cells and frozen at −80 for RNA extraction and library generation for RNAseq.

### *In vitro* Bone Marrow Derived Alveolar Macrophage culture

Femur, tibia and iliac bone were collected from CD11cCre+ *Klf4*^fl/fl^, and C57BL/6J animals. Bones were flushed with RPMI and red blood cells lysed with ACK buffer. Cells were counted and plated in 20ng/mL GM-CSF (Peprotech or R&D Systems) and 2ng/mL TGF-β (R&D Systems). On day 4, the cytokine-containing media on the cells was replaced. On day 8, cells were split and 0.1uM Rosaglitizone (Sigma) was added. Cells were harvested for analysis by flow cytometry on day 10.

### RNASeq library preparation, data mapping and analysis

Total RNA was isolated from cells using TRIzol or TRIzol LS reagent (Invitrogen). Chloroform extraction was performed according to the manufacturer’s protocol. RNA was recovered from the aqueous layer using RNA Clean and Concentrator-5 kit (Zymo Research). RNA was quantified with a Qubit (Invitrogen) and the integrity was evaluated with Agilent Bioanalyzer 2100 (Agilent Technologies). Libraries were prepared using the SMART-seq2 protocol (Picelli et al., 2014). Samples were sequenced on an Illumina NovaSeq sequencer using paired-end 50 base pair reads. Transcript abundances were quantified with the Ensembl GRCh38 cDNA reference using kallisto version 0.43.0. Transcript abundances were summarized to gene level using tximport. Differential expression statistics were generated using DEseq2. Genes with a Benjamini–Hochberg adjusted P-value <0.1 were called differentially expressed. GO term enrichment was performed with the DAVID Knowledgebase (v2023q3) web service.

### CUT&RUN and Library Prep

CUT&RUN was performed as previously described in (Skene, Henikoff and Henikoff, 2018) with modifications. Briefly, 600,000-800,000 Large Cavity Macrophages were sorted and each sample was split in half. Cells were then washed and bound to magnetic Con A beads (Bangs Laboratories) in 200uL PCR strip tubes. Cells were permeabilized with wash buffer containing 0.05% w/v Digitonin (Sigma-Aldrich). Each sample was incubated at 4°C with anti-GFP (Abcam, clone 290) or rabbit IgG (Sigma) at a concentration of 50μg/mL and left to rock overnight.

Permeabilized cells were washed and incubated rotating at room temperature for 10 min with pA-MNase (kindly provided by the Henikoff lab) at a concentration of 700ng/mL. After washing, cells were incubated at 0°C and MNase digestion was initiated by addition of CaCl2 to 1.3mM. After 2 hours, the reaction was stopped by the addition of EDTA and EGTA. Chromatin fragments were released by incubation at 37°C for 10 min. DNA was purified using DNA Clean&Concentrator-5 kit (Zymo). Libraries were prepared using the SMART-seq2 protocol (Picelli et al., 2014), quantified by Qubit (ThermoFisher), and quality checked by Bioanalyzer (Agilent) before sequencing on an Illumina NovaSeq as paired-ends to a depth of 3-5 million.

### CUT&RUN mapping, peak identification and motif analysis

Adapter sequence and low-quality bases were trimmed using TrimGalore (v0.6.7) and then mapped to the mouse reference genome (GRCm39) using Bowtie2 (v2.4.2). Peak analysis was performed using SEACR (v1.3) with corresponding merged control samples. The number of reads mapped to spike-in E. coli was used to normalize each sample and the top 1% of peaks are reported. To identify the motifs enriched in peak regions over the background, HOMER’s motif analysis (findMotifs.pl) including known default motifs and de novo motifs was used (Heinz et al., 2010). IgG control peaks were used as background.

### Statistical analysis

Data were analyzed with Prism software or algorithm (GraphPad Software). Statistical details and group sizes are indicated in the figure legends.

## Supplemental Materials

Figure 1 shows percentages and numbers of monocytes and neutrophils in *LysM*^+/+^, *LysM*^Cre/+^ *Klf2*^fl/fl^ and *LysM*^Cre/+^*Gata6*^fl/fl^ cavities, and gating schematics used for macrophages in various tissues throughout the paper. Figure 2 show the representative gating for the transferred macrophages and the RNASeq scatterplot with Gene IDs labeled. Figure 3 show the GFP expression in LCMs in the GFP-KLF2 mice, additional CUT&RUN peak figures for *MafB*, *Icam2*, *Timd4,* and GO term analysis for the differentially expressed genes from the RNASeq analysis of the *LysM*^+/+^*Klf4*^fl/fl^ vs *LysM*^Cre/+^*Klf4*^fl/fl^. Table S1 contains all antibodies, their clones, their manufacturers and the fluorophores used for flow cytometry.

## Data Availability

Sequencing results will be deposited to public databases. All other data are available in the main text or the supplementary materials. All materials (cell lines, mice, plasmids) are available upon request and after completion of a material transfer request to the corresponding author (GMB).

## Acknowledgements

We thank Dr. Jerry Lingrel for providing *LysM*^Cre^*Klf2*^fl/fl^ mice and Dr. Paul Kubes for providing *LysM*^Cre^*Gata6*^fl/fl^ mice. We thank the following core facilities at UC Berkeley: the Functional Genomics Laboratory (RRID:SCR_022170), the Vincent J. Coates Genome Science Laboratory (RRID:SCR_022170), and the Cancer Research Laboratory Flow Cytometry Facility. The CUT&RUN data analysis was performed by the Bioinformatics Core Facility at UC Davis.

We thank R. Vance, D. Raulet and members of the Barton and Vance labs for constructive discussions and critical reading of the manuscript. K.P. thanks Greg Timblin and Tessa Bergsbaken for their unwavering support. This work was supported by the NIH (R01AI072429 and R01AI158724 to G.M.B). G.M.B. is an Investigator of the Howard Hughes Medical Institute. K.P. was supported by the Cancer Research Institute as an Irvington Postdoctoral Fellow. This article is subject to HHMI’s Open Access to Publications policy. HHMI lab heads have previously granted a nonexclusive CC BY 4.0 license to the public and a sublicensable license to HHMI in their research articles. Pursuant to those licenses, the author-accepted manuscript of this article can be made freely available under a CC BY 4.0 license immediately upon publication.

## Disclosures

GMB is on the scientific advisory boards of X-Biotix and Actym Therapeutics and consults for Lycia Therapeutics. These activities are unrelated to the work described in this manuscript. The other authors have no disclosures and declare no competing interests.

## Author Contributions

K.P. and G.M.B. conceived of the project and wrote the manuscript. K.P. performed all experiments and analyzed all data. L.C.S. contributed to chimera analysis in Figures 1 and 5. All authors edited the manuscript.

## Abbreviations

AM: Alveolar Macrophage

ATRA: all-trans retinoic acid

LCM: Large Cavity Macrophage

OE: Overexpression

RV: Retrovirus

SCM: Small Cavity Macrophage

## Notes

### Competing Interest Statement

The authors have declared no competing interest.

