## Supplemental Figures 1 - 3 for "Krüppel-like Factor (KLF) family members control expression of genes required for serous cavity and alveolar macrophage identities"

Supplemental Figure 1- Pestal, et al

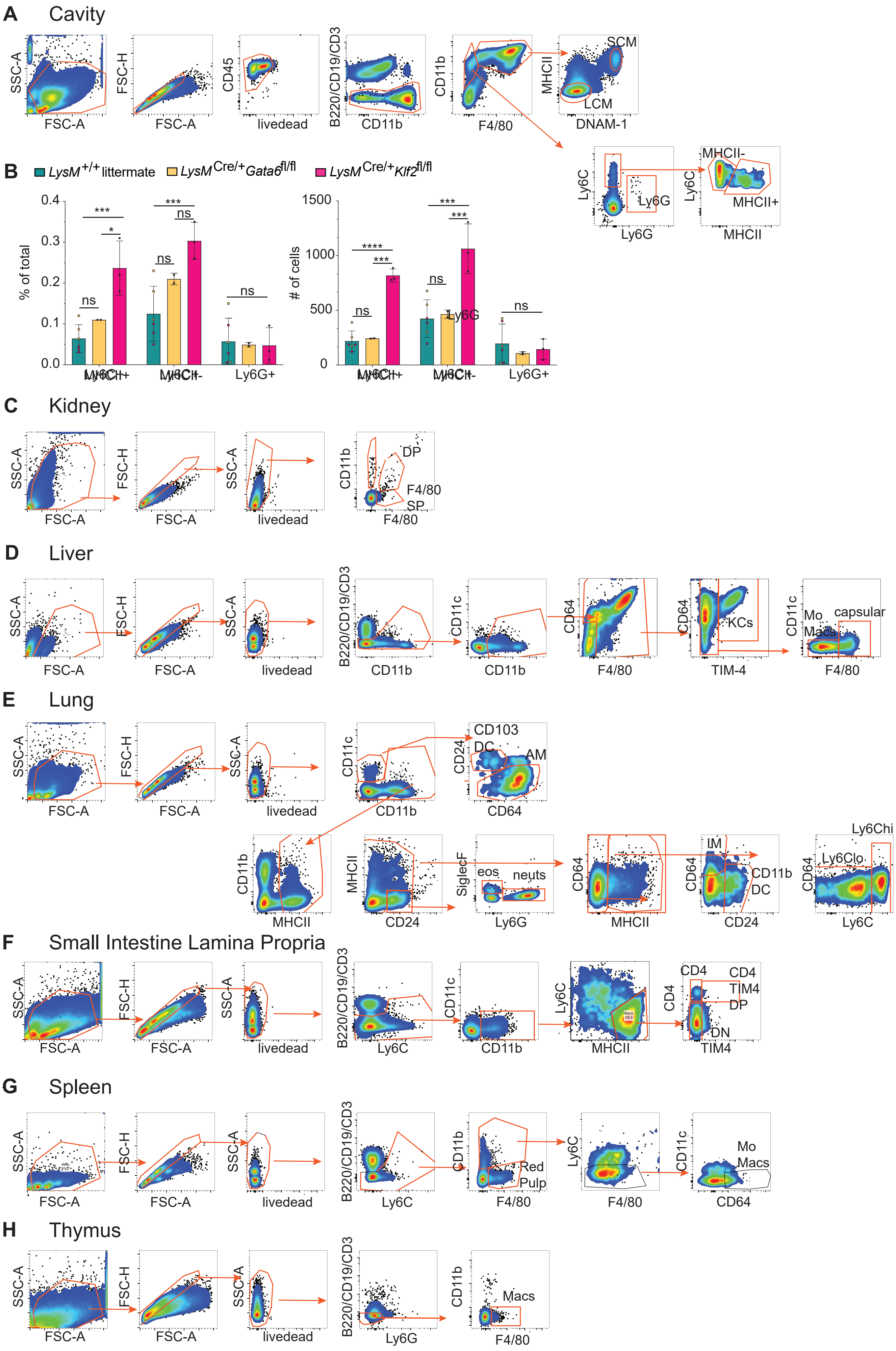

Fig S1

**A:** Flow cytometry gating strategy for cavities. **B:** Bar graphs depict the percent of total and total number of monocyte ($CD11b^+F4/80^-MHCII^+$ or $MHCII^-$) and neutrophils ($CD11b^+F4/80^-$, $Ly6G^+$), as measured by flow cytometry. Significance determined by ordinary 2-way ANOVA with multiple comparisons and Šidák's correction. Asterisks denote: ****<0.0001, *** 0.0006, ** 0.0021 , * 0.033). Flow cytometry gating for **C:** kidney, **D:** liver, **E:** lung, **F:** small intestine lamina propria, **G:** spleen, and **H:** thymus.

Supplemental Figure 2- Pestal, et al.

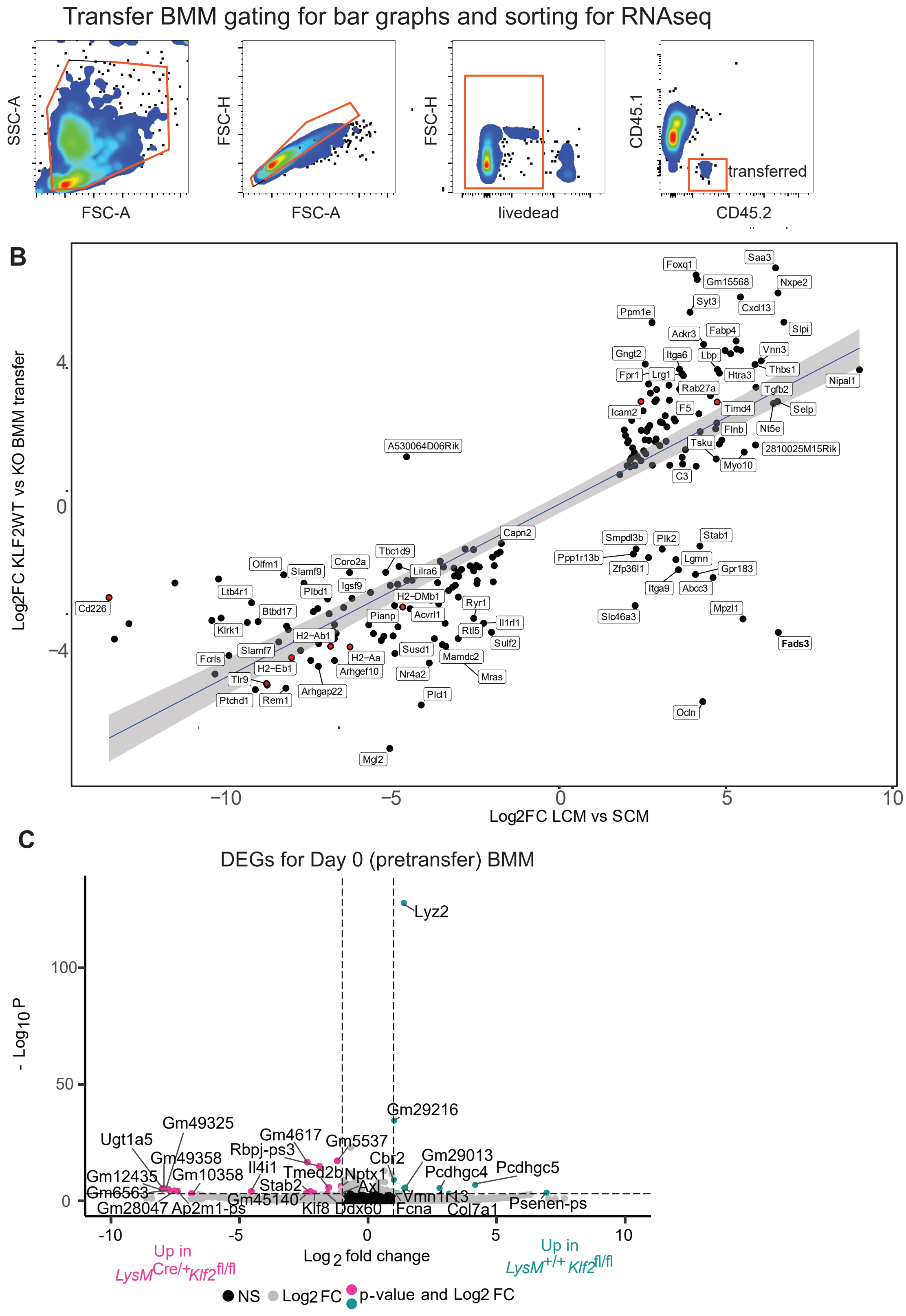


Fig S2

**A:** FACS gating for transferred bone marrow macrophages. **B:** Scatter plot of RNA Seq comparing differentially expressed genes between small and large cavity macrophages and transferred *LysM*$^{+/+}$*Klf2*$^{fl/fl}$ BMMs and *LysM*$^{Cre/+}$ BMMs with subset of genes labeled. **C:** Volcano plot of differentially expressed gene comparing pretransfer (day 0) *LysM*$^{+/+}$*Klf2*$^{fl/fl}$ to *LysM*$^{Cre/+}$*Klf2*$^{fl/fl}$ in vitro BMMs. Genes with a log2 Fold Change greater than |1| are colored grey. Genes with a p-value of -log10 (p value) greater than 4 are above the horizontal black line. Genes meeting both criteria are colored: teal, enriched in *LysM*$^{+/+}$*Klf2*$^{fl/fl}$; or pink, enriched in *LysM*$^{Cre/+}$*Klf2*$^{fl/fl}$.

Supplemental Figure 3- Pestal, et al

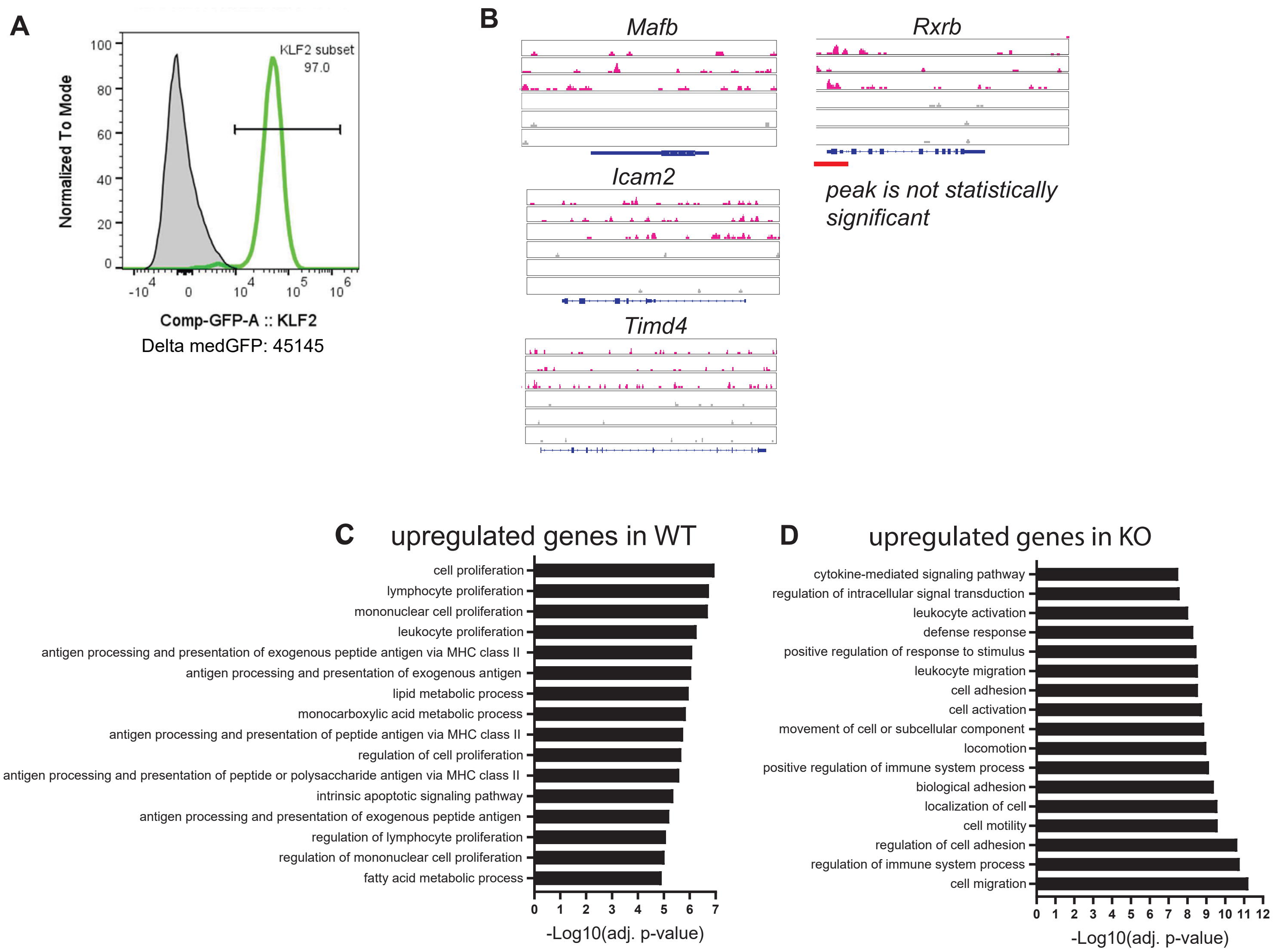


Fig S3

**A:** Representative flow cytometry of large cavity macrophages gated like SFig 1A (live, single, not B&T cells, $F4/80^{+}CD11b^{+}DNAM1^{-}MHCII^{-}ICAM2^{+}$) from GFP-KLF2 fusion positive mouse showing GFP expression. **B:** Genome browser tracks of anti-GPFKLF2 (pink) or IgG control (grey) CUT&RUN peaks from Large Cavity Macrophages in genes not directly regulated by KLF2 based on no peaks in the top 1% found by SEACR occur in two or more biological replicates. **C:** Top 25 GO term analysis results of genes upregulated in *LysM*$^{+/+}$*Klf4*$^{fl/fl}$ alveolar macrophages. **D:** Top 25 GO term analysis results of genes upregulated in *LysM*$^{Cre/+}$*Klf4*$^{fl/fl}$ alveolar macrophages

### Supplemental Table 1- Pestal, et al

| Marker | Clone(s) | Supplier:Fluorophore | | | | |
|---|---|---|---|---|---|---|
| CD8 | 53-6.7 | BD: BUV395 | | | | |
| I-A/I-E | M5/114.15.2 | BD: BUV496 | BioLegend: BV650 | | | |
| CD80 | 16-10A1 | BD: BUV563 | | | | |
| CD172a | P84 | BD: BUV615 | | | | |
| CD102 | 3C4(mIC2/4) | BD: BUV661 | BioLegend: AF647 | | | |
| CD209b | 22D1 | BD: BUV737 | | | | |
| CD11b | M1/70 | BD: BUV805 | BioLegend: BV785 | | | |
| Mer/MerTK | 108928 | BD: BV480 | | | | |
| H2 Class 1 | M1/42 | BD: BV510 | | | | |
| CD24 | M1/69 | BD: BV650 | Thermo: SuperBright 436 | | | |
| CD317 | 927 | Thermo: BV786 | | | | |
| Siglec-H | 440c | BD: BB700 | | | | |
| CD192 | 475301 | BD: RB780 | | | | |
| TIM-4 | 21H12; RMT4-54 | BD: R718 | BioLegend: PE-Cy7 | BioLegend: AF647 | | |
| CD19 | 6D5 | Bio-Rad: StarBright SBV440 | BioLegend: PE-Cy5 | | | |
| F4/80 | CI:A3-1; BM8 | Bio-Rad: StarBright SBB580 | BioLegend: PE | BioLegend: PE/Dazzle594 | | |
| CD3 | KT3; 145-2C11 | Bio-Rad: StarBright SBB615 | BD: PE-Cy5 | | | |
| NK1.1 | PK136 | Bio-Rad: StarBright SBB675 | | | | |
| Ly-6C | ER-MP20; HK1.4 | Bio-Rad: StarBright SBB765 | Thermo:APC-eF780 | BioLegend: AF700 | BioLegend: PE | Thermo: e450 |
| CD45RA/B220 | RA3-6B2 | Bio-Rad: StarBright SBB810 | BioLegend: PE-Cy5 | | | |
| CD163 | S15049I | BioLegend: BV421 | | | | |
| Ly-6G | 1A8 | BioLegend: Pacific Blue | BioLegend: AF700 | | | |
| CD4 | GK1.5 | BioLegend: Spark Violet 538 | Thermo: PE | | | |
| CD45 | 30-F11 | BioLegend: BV570 | | | | |
| XCR1 | ZET | BioLegend: BV605 | | | | |
| CD226 | TX42.1; 10E5 | BioLegend: BV711 | BioLegend: PE/Dazzle594 | | | |
| CD169 | 3D6.112 | BioLegend: PE/Dazzle 594 | | | | |
| CD274 | 10F.9G2 | BioLegend: PE/Fire 640 | | | | |
| CD86 | GL1 | BioLegend: PE-Cy5 | | | | |
| CD206 | C068C2 | BioLegend: PE/Fire 700 | | | | |
| CD64 | X54-5/7.1 | BioLegend: PE-Cy7 | | | | |
| CD103 | QA17A24 | BioLegend: PE-Fire 810 | | | | |
| CD11c | N418 | BioLegend: Spark NIR 685 | BioLegend: BV711 | | | |
| CD16/32 | S17011E | BioLegend: APC-Fire 750 | | | | |
| CX3CR1 | SA011F11 | BioLegend: APC/Fire 810 | | | | |
| MARCO | 579511 | R&D: APC | | | | |
| LYVE-1 | 223322 | R&D: Alexa Fluor 647 | | | | |
| Viability | | Thermo: LIVE DEAD Blue | Thermo: LIVE DEAD Aqua | | | |
| Siglec-F | 1RNM44N; E50-2440 | Thermo: PerCP-eFluor 710 | BD: AF647 | | | |
| CD45.2 | 104 | BioLegend: FITC | | | | |
| CD45.1 | A20 | Thermo: APC-eF780 | | | | |
| GATA6 | D61E4 | Cell Signaling Tech: PE | | | | |

Table S1

Antibodies used for flow cytometry. First row of fluorophores used for high-parameter spectral flow cytometry on a 5 laser Cytek Aurora. Other fluorophores were used with traditional flow cytometry on a BD LSR Fortessa.
